## Supplementary Data for "Coupling chromosome organization to genome segregation in Archaea"

### Supplementary Figures

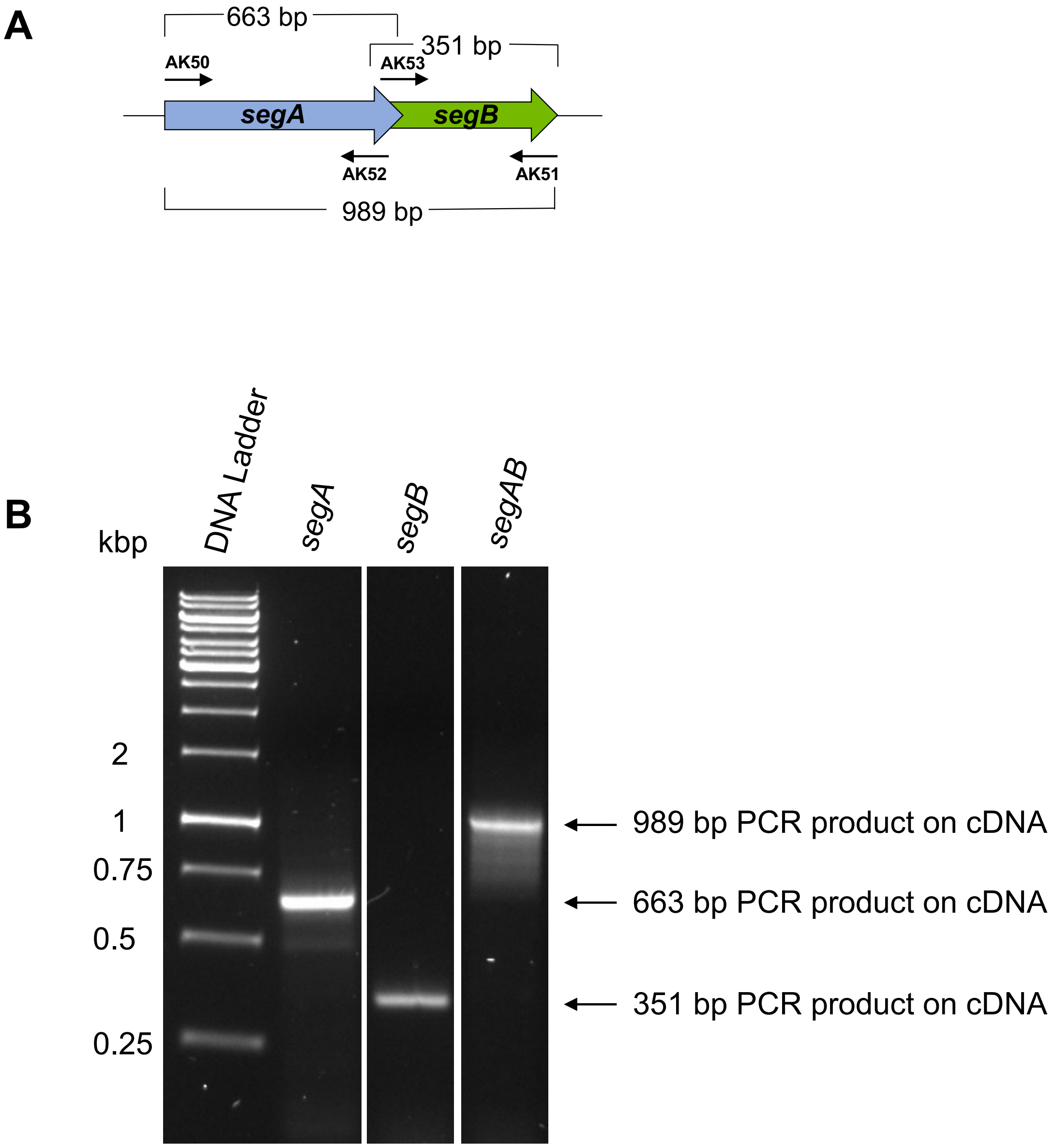


**Supplementary Figure 1. The *segAB* genes form an operon. (A)** Schematic of the *segAB* locus in *S. solfataricus*. Primers used to amplify fragments are indicated (not to scale) and the predicted amplicon sizes are shown. **(B)** PCR products obtained using cDNA as template and analysed on a 1% agarose gel stained with Syber Safe.

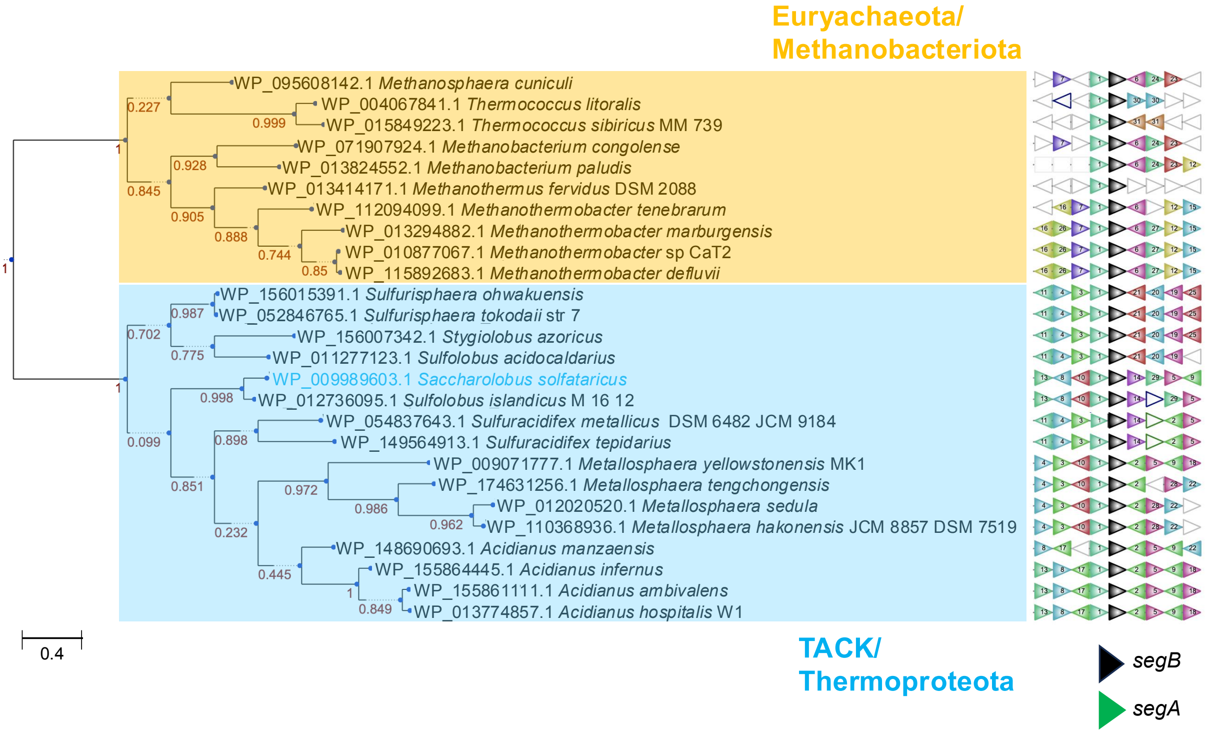


**Supplementary Figure 2. Phylogenetic tree of SegB orthologues among archaea.** The tree was constructed using the FlaGs programme[1] that deploys the tree-building feature of ETE v3 Python environment[2]. Bootstrap values are shown next to the branches. The two major clades are highlighted in yellow-orange (Euryarchaeota/Methanobacteriota) and blue (TACK/Thermoproteota). The genomic neighbours of each *segB* orthologue are shown on the diagram on the right side, in which the arrows represent genes that are coloured and numbered in the same way as in Figure 1. The *segB* gene is shown in black and *segA* gene in green.

***Metallosphaera hakonensis* DSM7519 (300 aa) (12.5 repeats)**

MSEIDFLLKRKKKNEGVQGGESGAQRVESRVQGGESGAQRVESRVQGGESGAQRVESRVQGGESGAQRVESRVQG
GESGAQRVESRVQGGESGAQRVESRVQGGESGAQRVESRVQGGESGAQRVESRVQGGESGAQRVESRVQGGESGA
QRVESRVQGGESGAQRVESRVQGGESGAQRVESRVQGGESGAQTTIDITLDSTGIVRTPESRAVSYVGTGIAQES
SSVNLDLMENFLDKDPKIGVWSYPSYIVLQYLYHVKPGFKMSKIAKEALELGMRQLFPDLYSKAEELSRKKGLLK

***Metallosphaera sedula* (272 aa) (11 repeats)**

MSELDFLLKRKKKTEGVQGGESGGQRVESRVQGGESGGQRVESRVQGGESGGQRVESRVQGGESGGQRVESRVQG
GESGGQRVESRVQGGESGGQRVESRVQGGESGGQRVESRVQGGESGGQRVESRVQGGESGGQRVESRVQGGESGG
QRVESRVQGGESGGQRVESIQISPETDSHQNGNTEQTTQTVTQTSVDQTSLEQLMESLLDKDPKIGVWSYPGYAV
LQYLYHTKPGFRMSKVAKDALELGLKQLYPDLYSKAESLARQKGLIK

***Metallosphaera tengchongensis* (247 aa) (8.5 repeats)**

MSELDFLLKRKKKDEQKSGESVEKRTEGRVQSGESVEKRTEGRVQSGESVEKRTEGRVQSGESVEKRTEGRVQSG
ESVEKRTEGRVQSGESVEKRTEGRVQSGESVEKRTEGRVQSGESVEKRTEGRVQSGESVERPNIHEVELNVVTIG
NVKPDLPPRDDAQENTETPSSGIDEVEKLMGNFLDKDPKIGVWSYPSYMVLQYLFHTKPGFKMSKMAKDALEIGM
RQLFPDLYSKAEKIAKDKGLLR

***Metallosphaera cuprina* (269 aa) (11 repeats)**

MSELDFLLNRKKRDGGVQGVDNSPQRVDNRVQGVDNSPQRVDNRVQGVDNSPQRVDNRVQGVDNSPQRVDNRVQG
VDNSPQRVDNRVQGVDNSPQRVDNRVQGVDNSPQRVDNRVQGVDNSPQRVDNRVQGVDNSPQRVDNRVQGVDNSP
QRVDNRVQGVDNSPQRVDNTIDSGADKSPMISLENQNVKRDQRLEDEELELMRKFLDKDPKIGIWSYPSYVVLQY
LFNTRPGFKISKVAKEALEIGMRQLFPDLYTKAELITKQRGLVK

***Metallosphaera yellowstonensis* MK1 (172 aa) (no repeats)**

MSELDFLLNRKKSVGKTSDSRVEKEEEIGLESRENRGEALPQTTESSGKSVEERGKPEESSGLTLESLGKSVEGT
PEVEMETLEERGNPLQSGVSRDKRSIESVMMTLLSREPKIGVWSYPSFLVLQYLFSTKPGFRMSKIAKEALEIGL
RQLYPELFAIAESVAKEKGLIK

***Methanobacterium congolense* (168 aa) (no repeats)**

MSQKKKRESALGRGLDALIRTQPVEEPGEKEQDVESEIPAEEEKPTQRKTPPAKKTRKPSKTSGRSTAKTARKSP
EKAQKPKIPRPKKKPETAAEDFNVDSQLLEEVMAEVAKNPRISLWSAKSAAVLRYLRKTKPAFSISKEASALIED
AVKEKYPDLWEIFEGEGL

***Acidianus manzaensis* (170 aa) (no repeats)**

MSELDFLLNRKKNLEKKDKKEESREETKTNTEEKIQNTQEQIEKSEERKTEETKIQNIDSIENSSNSIVMKEESA
ENNQYSREEKTENSRVENAEDREDNVESIMKTFLNKDPKIGVWSYPSFLVLQYLYNTKPGFKMSKVAKDALEYGL
KRMYPELFDKAEKISQSKIR

***Sulfolobus acidocaldarius* (151 aa) (2 short repeats)**

MSELDLILNRKKTEQKVEASEQKVESRQETSKSTTQKLAETKTSETQEKVNENDASKSPREDLEEVKSSPNSNSE
VKVDNIKGIMKKFIDRDPKIGIWSYPSFLVLQYLYHTVPGFKMSKIAKDALERGLREIYPDLFRIAEEVTLESKK
S

***Saccharolobus solfataricus* (109 aa) (no repeats)**

MSELDFLLKKKRKSEDEEKIINNNENAKKEEITNEEEKIKNDMLKYIEKDPKIGVWSYPAFLVLQYLYHTVPGFK
MSRTAKEALEKGLKEMYPTLFTIAEKIAKERFKE

**Supplementary Figure 3. SegB orthologues harbour repeat sequences.** Amino acid sequences of SegB proteins in different archaeal genera. The repeats are highlighted in colour.

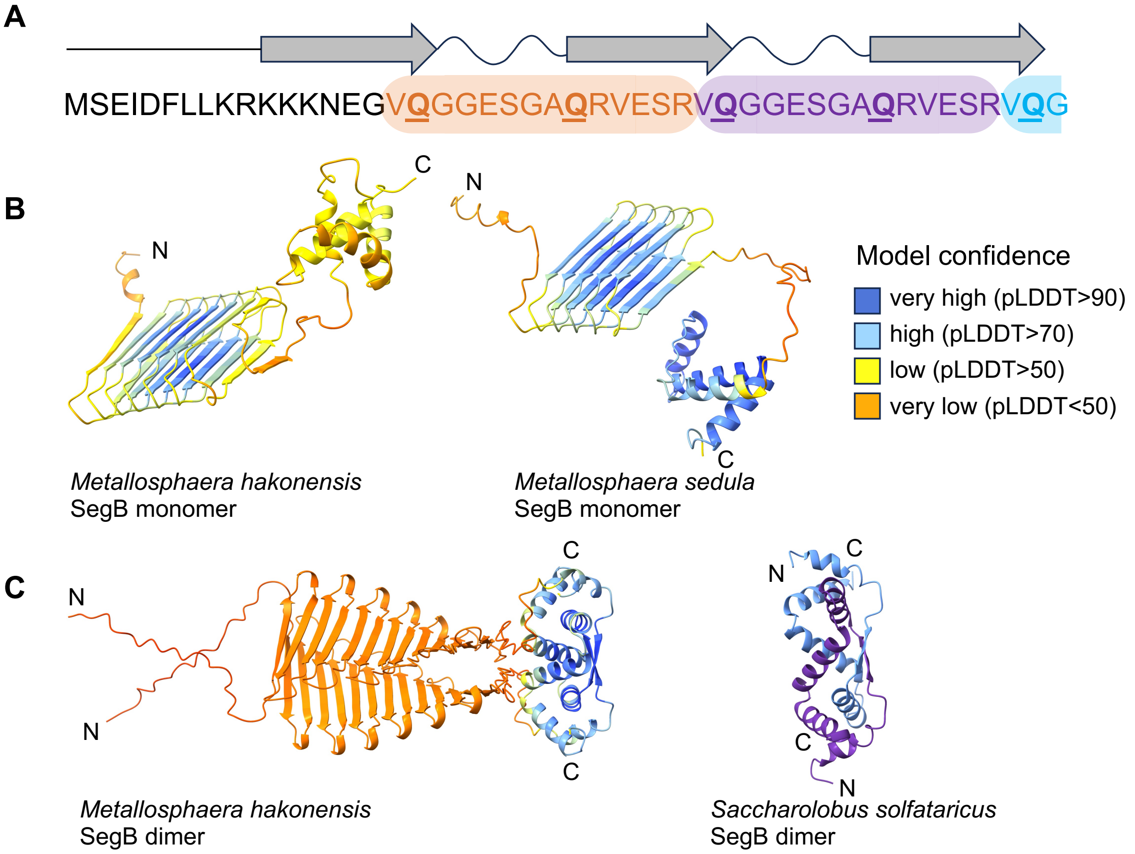


**Supplementary Figure 4. SegB from *Metallosphaera hakonensis* contains repeats and has a predicted β-helix structure domain. (A)** SegB protein sequence in which the repeats are highlighted in coloured boxes and the predicted secondary structure is shown above. Each repeat starts at the end of a β-strand and extends until the next β-strand, including the intervening loop. **(B)** AlphaFold2-predicted structures of *Metallosphaera hakonensis* and *Metallosphaera sedula* SegB monomer, coloured according to level of confidence. **(C)** AlphaFold2-predicted structure of *Metallosphaera hakonensis* SegB dimer (left) and experimentally determined X-ray structure of *S. solfataricus* SegB dimer (right).

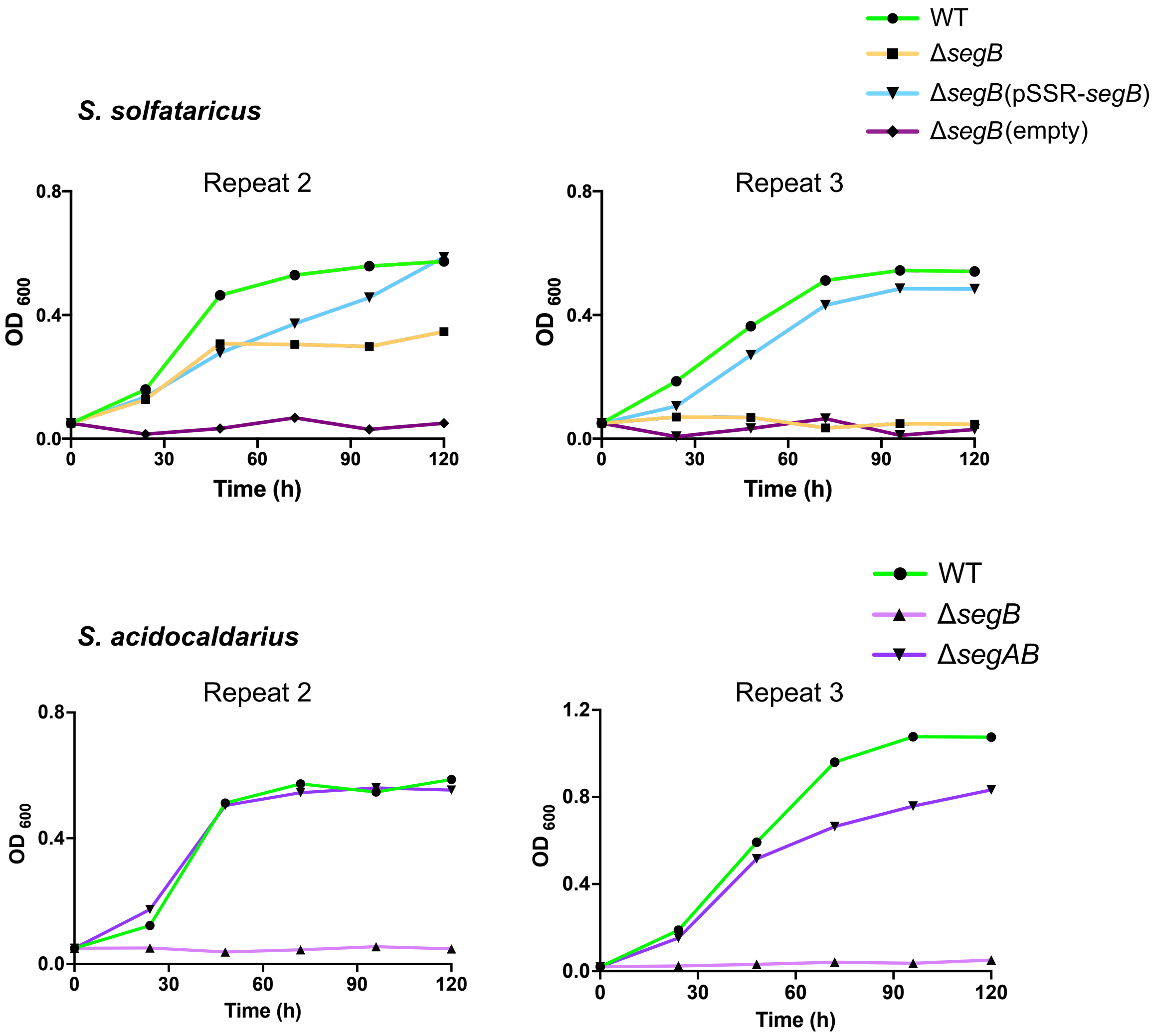


**Supplementary Figure 5. Repeats of growth experiments.** Cultures were grown at 75°C in Brock's medium supplemented with either 0.1% tryptone or NZ-amine, 0.2% sucrose and uracil (20 μg/mL), when necessary. *S. solfataricus* strains that carry the simvastatin-resistance selection marker were grown in Brock's medium supplemented with simvastatin (15 - 20 μM). Repeats of *S. solfataricus* **(A)** and *S. acidocaldarius* **(B)** growth curves.

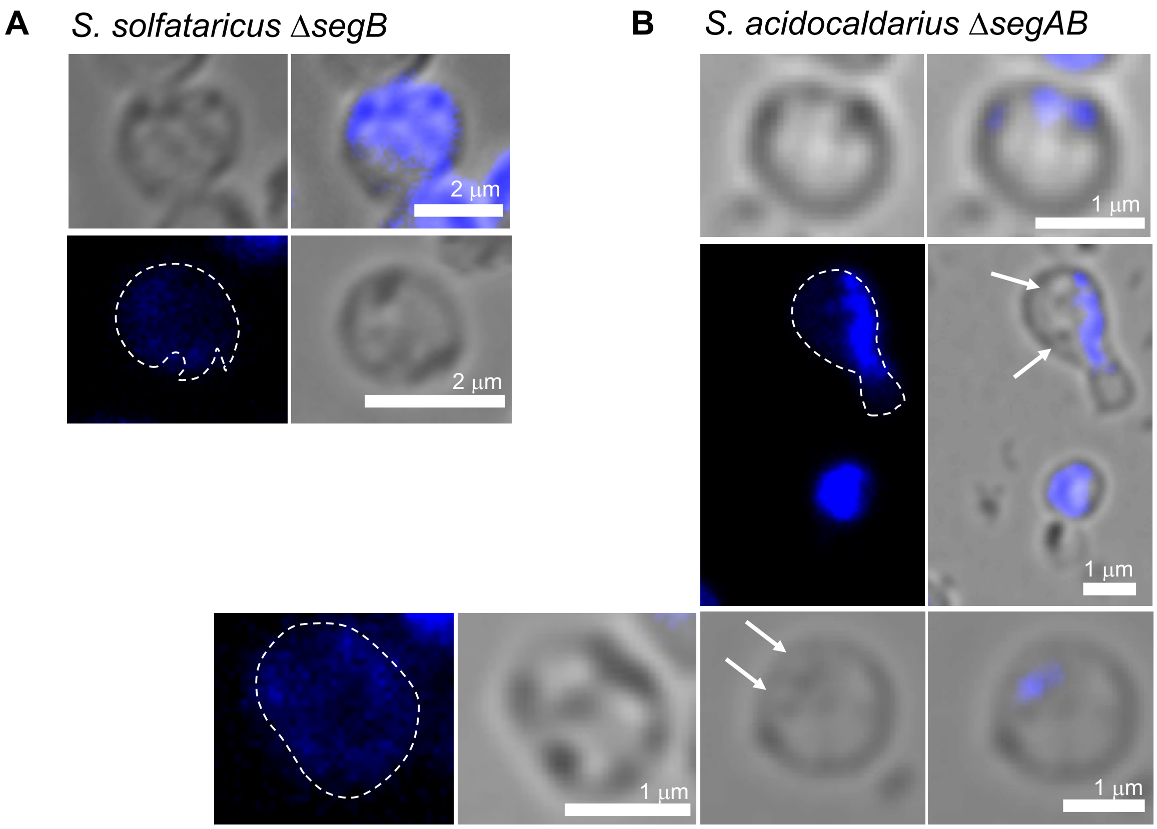


**Supplementary Figure 6. Deletion strains exhibit different aberrant phenotypes.** Microscopy images of DAPI-stained cells of *S. solfataricus* **(A)** and *S. acidocaldarius* **(B)** deletion strains. The white arrows point to blebs. The scale bar is either 1 or 2 μm.

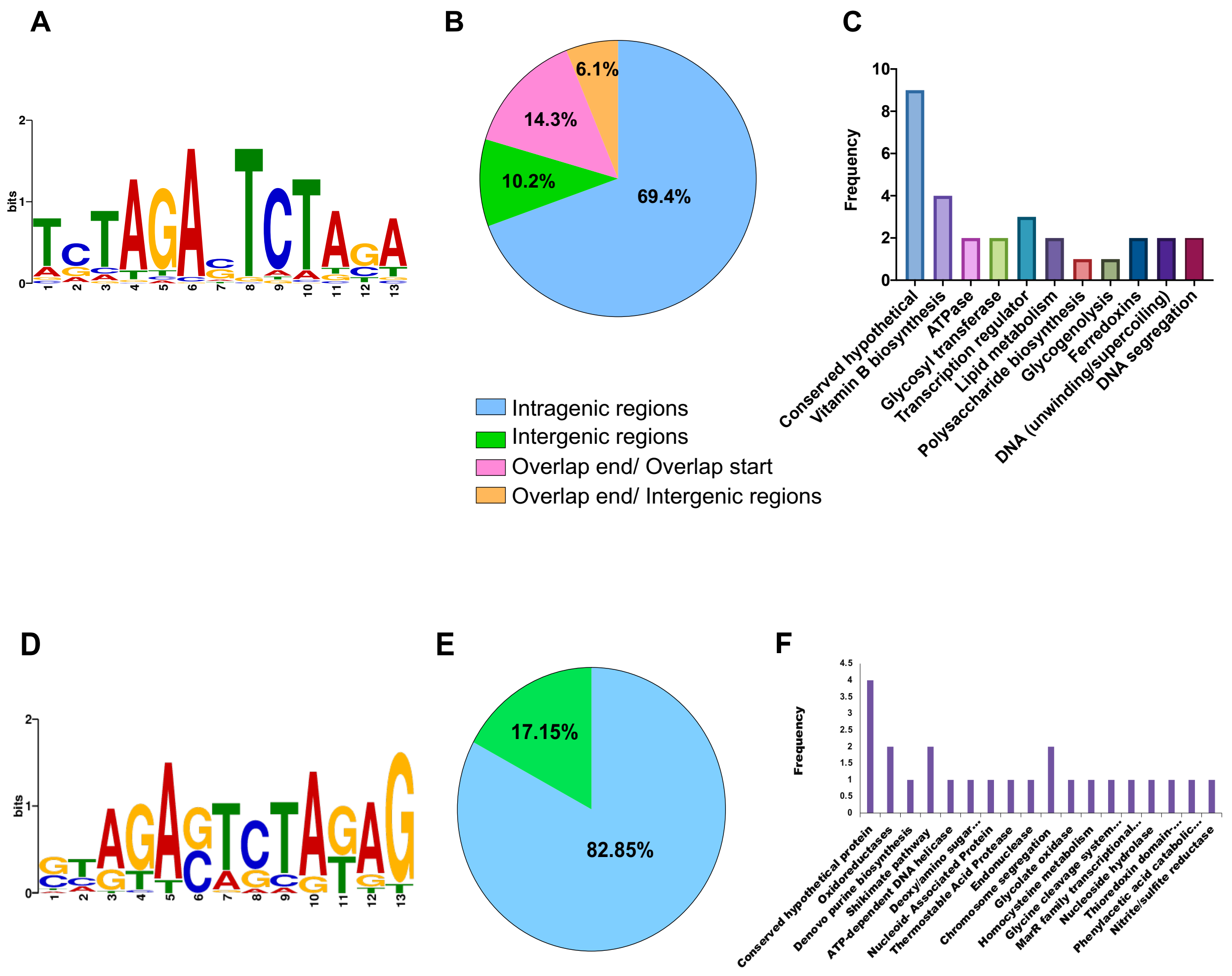


**Supplementary Figure 7. SegB binds to multiple sites on the chromosome of *S. solfataricus* and *S. acidocaldarius*. (A)** DNA-binding consensus motif for *S. solfataricus* SegB identified by MEME-ChIP (with an E-value of 8.3e$^{-}$ 039) based on the 49 sites present in 39 enrichment peaks. **(B)** Pie chart showing the percentages of *S. solfataricus* SegB sites in intragenic or intergenic regions, or relative to the closest gene. **(C)** Established or hypothetical function of proteins encoded by genes that are in the intragenic category in panel B. **(D)** DNA-binding consensus motif for *S. acidocaldarius* SegB identified by MEME-ChIP (with an E-value of 1.7e-019) based on the 35 sites present in 32 enrichment peaks. **(E)** Pie chart showing the distribution of *S. acidocaldarius* SegB sites in either intragenic or intergenic regions. **(F)** Established or hypothetical function of proteins encoded by genes that are in the intragenic category in panel E.

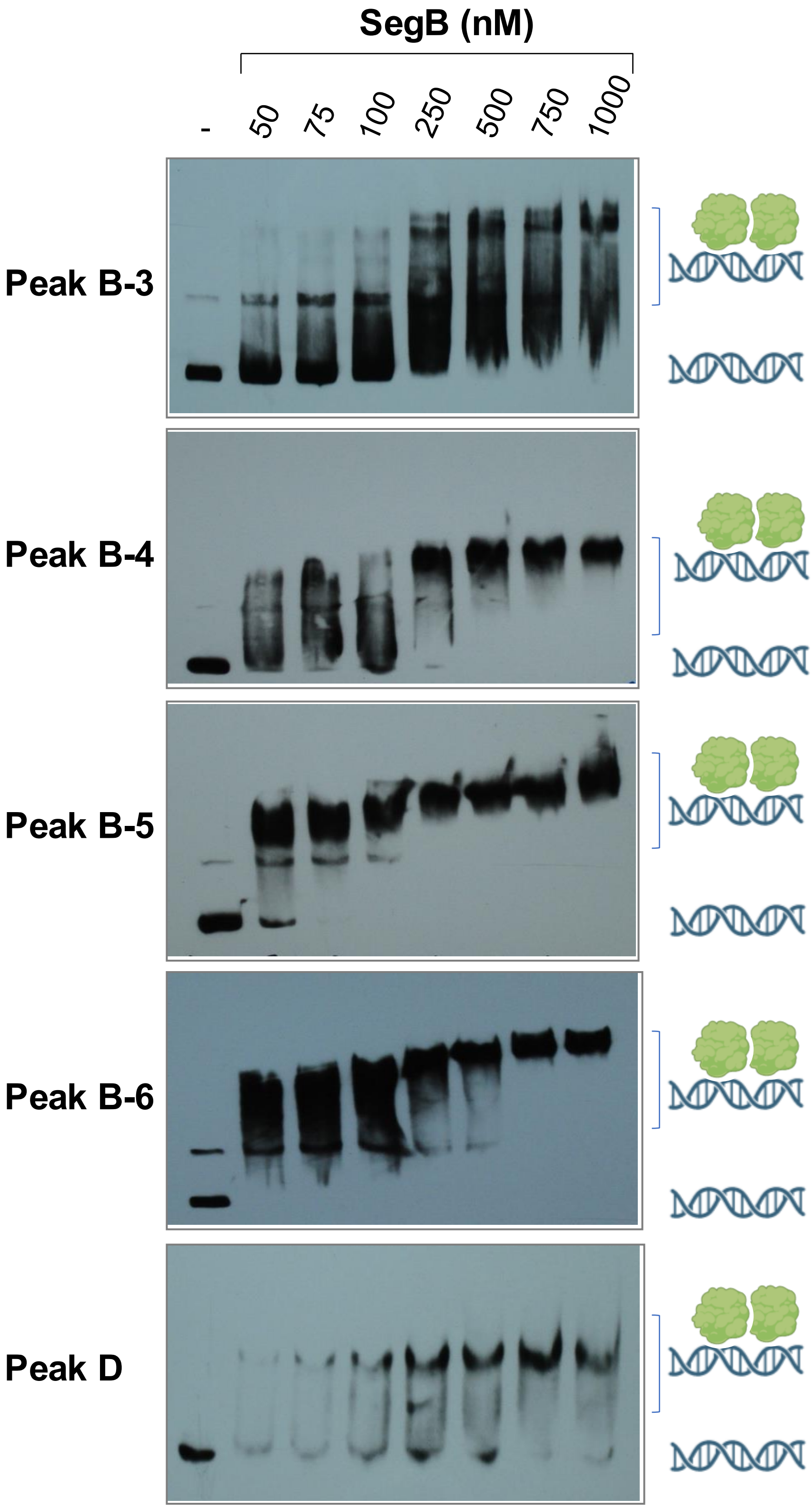


**Supplementary Figure 8. *In vitro* validation of the SegB binding sites identified by ChIP-seq.** EMSAs in which biotinylated DNA fragments (1-5 nM) harbouring the sequence corresponding to different enrichment peaks were incubated with *S. solfataricus* SegB in the presence of competitor polydIdC DNA (1 μg). Unbound DNA and complexes are indicated by the cartoon images.

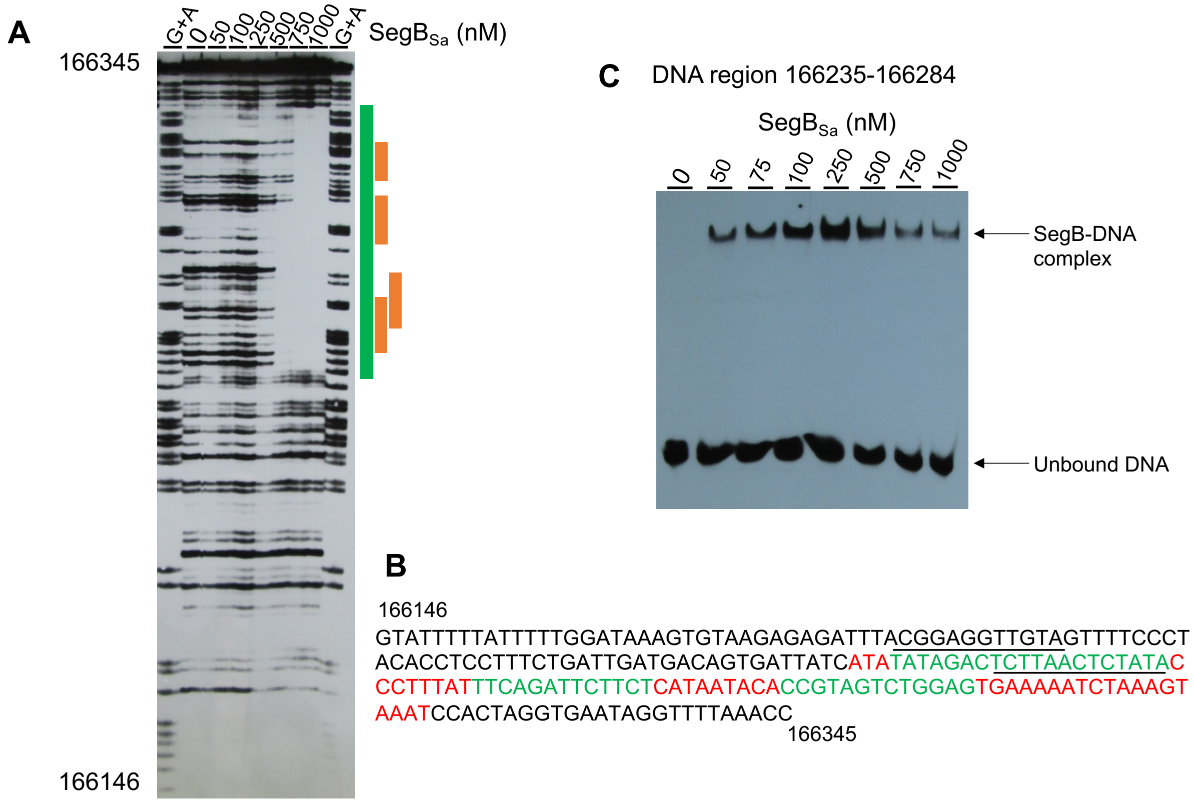


**Supplementary Figure 9. *S. acidocaldarius* SegB binds to the DNA region upstream of the *segAB* locus. (A)** DNase I footprint performed with a biotinylated DNA fragment spanning the region upstream of the *segAB* genes, including part of the *segA* gene, and increasing concentrations of SegB. The window of protection is indicated by the green bar and the potential binding sites are denoted by the orange bars. **(B)** Sequence of the DNA fragment deployed for the DNase I footprint (genomic coordinates: 166146-166345). The sequence highlighted in red corresponds to the window of protection and the sequences shown in green are the potential SegB binding sites. The sequence closer to the 166146 bp end contains two potential overlapping sites (TATAGACTCTTAA and TCTTAACTCTATA), indicated with a line above and below the text, respectively. **(C)** EMSA in which increasing SegB concentrations were incubated a biotinylated oligonucleotide spanning the region upstream of *segA*.

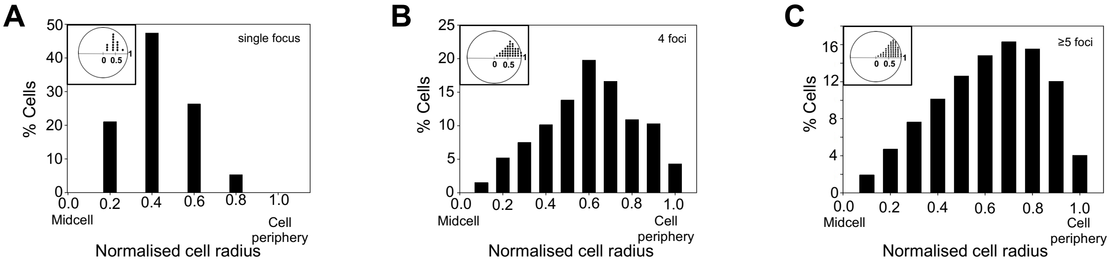


**Supplementary Figure 10. Distribution of SegB foci along the cell radius: foci number increase correlates with a shift of the prevalent position towards the cell membrane.** Analysis of the position of SegB foci in cells containing a single focus **(A)** (n=19), four foci **(B)** (n=196) and five or more foci **(C)** (n=557). The inset in the top left corner is a diagram indicating the location of the foci within the cell.

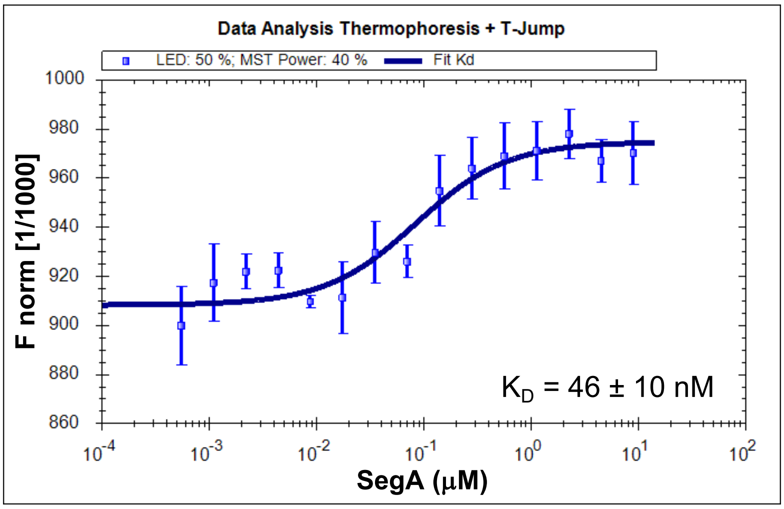


**Supplementary Figure 11. SegA interacts with SegB with high affinity.** Microscale thermophoresis (MST) experiments were performed mixing NT-647-labelled SegB (304 nM) with increasing concentrations of SegA (0.6, 1.2, 2.4, 4.9, 9.8, 19.5, 39, 78, 156, 312, 625, 1250, 2500, 5000 and 10000 nM). measurements were carried out using the following parameters: 20% LED power, 40% MST power, 5 second delay before heating, 30 seconds during MST, 5 second delay after heating, 15 second delay. Binding data were analysed using NTAnalysis software (NanoTemper Technologies). The average values from three replicates were used to plot the binding curve and derive error.

### Supplementary Tables

**Supplementary Table Primers. Oligonucleotides used in this study.**

| Primer name | 5'–3' sequence | Details |
|---|---|---|
| segBKOUSforward Kpn | CCCGGTACCGCATATAAATATTATCA TTATAATATGGTGGACTCG | Forward primer to clone the 823 bp region upstream of *S. solfataricus segB* into pET2268 for knock out construction |
| segBKOUSreverse Nco | CCCCCATGGTTTCTTTTCTTCTTCAA T | Reverse primer to clone the 823 bp region upstream of *S. solfataricus segB* into pET2268 for knock out construction |
| segBKODSforward Bam | CCCGGATCCCCCTACACTATTTACC ATTGCAGAAAAAATTGC | Forward primer to clone the 1150 bp region downstream of *S. solfataricus segB* into pET2268 for knock out construction |
| segBKODSreverse Not | CCCGCGGCCGCCCCTTTTTCATGTT AAAAATTCACG | Reverse primer to clone the 1150 bp region downstream of *S. solfataricus segB* into pET2268 for knock out construction |
| saci0203 pyrEF for | AGCTAAAGTGGTGATAGAAGGATGA GTGAGTTAGACTTAATCTTAAACAGT TTGAGCAGTTCTAG | Forward primer for PCR to knock out *S. acidocaldarius segB* (*saci_0203*) gene by insertion of *pyrEF* cassette |
| saci0203 pyrEF rev | CCCTACTTTGGTAATATTCATTATAG TAATATCATTAACTCTTTTTACTCGA CCGGCTATTTTTTCAC | Reverse primer for PCR to knock out *S. acidocaldarius segB* and *segA* (*saci_0203* and *saci_0204*) genes by insertion of *pyrEF* cassette |
| saci0203_saci0204 pyrEF for | GAATCTGAAATAAAGGGTATAGAGT TAAGAGTCTATATATGATAATCACTG TTTGAGCAGTTCTAG | Forward primer for PCR to knock out *S. acidocaldarius segB* and *segA* (*saci_0203* and *saci_0204*) genes by insertion of *pyrEF* cassette |
| saci0204(-47) FP | BTN- TTATGAGAAGAATCTGAAATAAAGG GTATAGAGTTAAGAGTCTATATATG | Biotinylated forward oligonucleotide containing the sequence 50 bp upstream of *S. acidocaldarius segB* (*saci_0204*) start codon used for EMSA |
| saci0204(-47) RP | CATATATAGACTCTTAACTCTATACC CTTTATTTCAGATTCTTCTCATAA | Reverse oligonucleotide containing the sequence 50 |

| | | |
|---|---|---|
| | | bp upstream of *S. acidocaldarius segB* (*saci_0204*) start codon used for EMSA |
| segA-stop-Rev | CTCGAGTCATTCACTAATCACCTTTGCTAATTGC | Reverse primer containing stop codon for PCR amplification of *segA* and cloning into pET22b (XhoI) |
| AK50 | ATGATAGTCACAGTAATAAATCAGAAAG | Forward primer to amplify *segA* from *S. solfataricus* P2 for RT-PCR |
| AK51 | TTACTCTTTAAATCTCTCTTTTGC | Reverse primer to amplify *segB* from *S. solfataricus* P2 for RT-PCR |
| AK52 | TCATTCACTAATCACCTTTGCTAATTG | Reverse primer to amplify *segA* from *S. solfataricus* P2 for RT-PCR |
| AK53 | ATGAGTGAATTAGATTTCCTATTG | Forward primer to amplify *segB* from *S. solfataricus* P2 for RT-PCR |
| AK80 | ATACATATGAGTGAATTAGATTTCC | Forward primer to amplify *segB* from *S. solfataricus* P2 (NdeI) for cloning into pSSR vector |
| AK81 | ATAATAATCGATTTACTCTTTAAATCTCTC | Reverse primer to amplify *segB* from *S. solfataricus* P2 (ClaI) for cloning into pSSR vector |
| AK87 | ACGCTATGTAGGCTGAAGCTA | Forward primer to amplify a 289 bp DNA fragment from *S. solfataricus* P2 for AFM experiments |
| AK97 | AGTAAGATAAATTTACTGAAGTCG | Reverse primer to amplify a 289 bp DNA fragment from *S. solfataricus* P2 for AFM experiments |
| AK93 | ATAATAGCATGCAAGGGAAGTAGAG | Forward primer to amplify a 1028 bp DNA fragment (containing peak D) from *S. solfataricus* P2 (SphI) for cloning into pUC18. The plasmid was used in AFM experiments. |
| AK94 | ATAATAGGATCCAGACTGGCTAAGAC | Reverse primer to amplify a 1028 bp DNA fragment |

| | | |
|---|---|---|
| | | (containing peak D) from *S. solfataricus* P2 (BamHI) for cloning into pUC18. The plasmid was used in AFM experiments. |
| AK95 | ATAATA<u>GGATCC</u>AAGGGAAGTAGAG | Forward primer to amplify a106 bp DNA fragment (peak D) from *S. solfataricus* P2 (BamHI) for cloning into pNER803. The plasmid was used in AFM experiments. |
| AK96 | ATAATA<u>GGTACC</u>TAATTAATAGTTAACAC | Reverse primer to amplify a106 bp DNA fragment (peak D) from *S. solfataricus* P2 (KpnI) for cloning into pNER803. The plasmid was used in AFM experiments. |
| NER36 | ATAATA<u>AAGCTT</u>AAGGAAGTAGAGTC | Forward primer to amplify a 1028 bp DNA fragment (containing peak D) from *S. solfataricus* P2 (HindIII) for cloning into pNER804. The plasmid was used in AFM experiments. |
| NER37 | ATAATA<u>GCATGC</u>AGACTGGCTAAGAC | Reverse primer to amplify a 1028 bp DNA fragment (containing peak D) from *S. solfataricus* P2 (SphI) for cloning into pNER804. The plasmid was used in AFM experiments. |
| NER7 | TTTAAATGTATATCCTCGC | Forward primer to amplify *S. solfataricus* P2 genomic region 20475-20674 (peak B4) for EMSA and DNaseI footprint |
| NER8 | AGGGTAATGATATGGACTT | Reverse primer to amplify *S. solfataricus* P2 genomic region 20475-20674 (peak B4) for EMSA and DNaseI footprint |
| NER9 | GATTTTTATGAATTAGGATTCA | Forward primer to amplify *S. solfataricus* P2 genomic region 22703-22902 (peak B5) for EMSA and DNaseI footprint |

| | | |
|---|---|---|
| NER10 | GCATCTATGGCATCTG | Reverse primer to amplify *S. solfataricus* P2 genomic region 22703-22902 (peak B5) for EMSA and DNaseI footprint |
| NER11 | GCTCGTAATTGAGGC | Forward primer to amplify *S. solfataricus* P2 genomic region 24311-24510 (peak B6) for EMSA and DNaseI footprint |
| NER12 | ACTTAACTCTAGATAATAAATC | Reverse primer to amplify *S. solfataricus* P2 genomic region 24311-24510 (peak B6) for EMSA and DNaseI footprint |
| NER47 | CTCAGCTGTCGGTAATAC | Forward primer to amplify *S. solfataricus* P2 genomic region 18531-18730 (peak B3) for EMSA and DNaseI footprint |
| NER48 | AGGAGTATGGAATACGTC | Reverse primer to amplify *S. solfataricus* P2 genomic region 18531-18730 (peak B3) for EMSA and DNaseI footprint |
| AK85-B-forw | GAATGTCACAAGGTTCTT | Forward primer to amplify *S. solfataricus* P2 genomic region 1001932-1002131 (peak D) for EMSA and DNaseI footprint |
| AK86-B-rev | AACACTCTAACTATTTATG | Reverse primer to amplify *S. solfataricus* P2 genomic region 1001932-1002131 (peak D) for EMSA and DNaseI footprint |
| NER42 | GTATTTTTATTTTTGGATAA | Forward primer to amplify *S. acidocaldarius* genomic region 166146-166345 for DNaseI footprint |
| NER43 | GGTTTAAAACCTATTCAC | Reverse primer to amplify *S. acidocaldarius* genomic region 166146-166345 for DNaseI footprint |
| Saci_0203_Forw | ATATATCATATGAGTGAGTTAGACTTAGAC | Forward primer to amplify *S. acidocaldarius segB* gene |

| | | |
|---|---|---|
| | | (*saci_0203*) for cloning into pET22b (NdeI) |
| Saci_0203_Back | ATATATCTCGAGACTCTTTTTACTCTCTAA | Reverse primer to amplify *S. acidocaldarius segB* gene (*saci_0203*) for cloning into pET22b (XhoI) |

**Supplementary Table Plasmids. Constructs used in this study.**

| Plasmid | Description | Source/Reference |
|---|---|---|
| pET22b | Expression vector containing the T7 promoter and the sequence encoding a hexa-his-tag at the 3' of the cloned gene | Novagen |
| pET-*sso035* | Expression plasmid containing the *S. solfataricus segB* gene with no STOP codon cloned into NdeI and XhoI restriction sites | Kalliomaa-Sanford *et al* (2012)[3] |
| pET22b-*sso034*-stop | Expression plasmid containing the *S. solfataricus segA* gene with STOP codon cloned into NdeI and XhoI restriction sites | This work |
| pET22b-*saci0203* | Expression plasmid containing the *S. acidocaldarius segB* (*saci_0203*) gene with no STOP codon cloned into NdeI and XhoI restriction sites | This work |
| pET2268 | Plasmid used for the construction of *S. solfataricus* Δ*segB* strain and containing *lacS* gene | Albers and Driessen (2008)[4] |
| pET2268-segB-KO-US-fragment | pET2268 containing the 823 bp region upstream of *S. solfataricus segB* cloned into KpnI and NcoI restriction sites upstream of the *lacS* gene | This work |
| pET2268-segB-KO-DS-fragment | pET2268 containing the 1150 bp region downstream of *S. solfataricus segB* cloned into BamHI and NotI restriction sites downstream of the *lacS* gene | This work |
| pSVA406 | pGEM-T derived plasmid containing the *pyrEF* cassette from *S. solfataricus*, used for overlap extension PCR to construct *S. acidocaldarius* deletion strains | Wagner *et al* (2012)[5] |
| pSSR | Expression vector for *S. solfataricus* | Zheng *et al* (2012)[6] |
| p*segB* | pSSR containing *S. solfataricus segB* gene | This work |
| pUC18 | Cloning vector | Norrander *et al* (1983)[7] |
| pNER803 | pUC18 containing a 1028 bp *S. solfataricus* region that harbours peak D cloned into SphI and BamHI restriction sites | This work |
| pNER804 | pNER803 containing a 126 bp *S. solfataricus* region that harbour the high-affinity SegB binding site located in peak D, cloned into BamHI and KpnI restriction sites | This work |

| | | |
|---|---|---|
| pNER805 | pNER804 containing a 1028 bp *S. solfataricus* region that harbours peak D cloned into SphI and HindIII restriction sites | This work |
