## Supplementary Table S1 for "Coupling chromosome organization to genome segregation in Archaea"

| #Peak ID-Figure 4 | Start | End | Strand | Position of the peak | Gene accession number/name | Product (gene function) | DNA sequence under peak<br>Motifs identified by MEME (yellow)<br>Further motifs identified by DNaseI footprint (blue) |
|---|---|---|---|---|---|---|---|
| B-1 | 1576 | 1748 | + | intragenic | SSO_RS00015 (SSO0002)(*thiD*) | Phosphomethylpyrimidine kinase (Thiamine biosynthesis) | AAATTCGCTACAATGACAGTCAAATACGG<br>TTTGGACTTAGGAGGAGGATATGGACCAG<br>TAGATCCCTTTGCCCCTATAGAGTCCATA<br>GTGAAGAGAGAAGAAGGAAGAAATCAGCT<br>AGAAAACTTACTTTGGTACTTAGAGTCTA<br>ATCTTAACGTTATACTTAAACTAATTAA |
| B-2 | 14998 | 15178 | + | intragenic | SSO_RS00080 (SSO0016) (*queC* or *exsB like*) | ExsB transcription regulator (Queuosine biosynthesis) | ACAATATATCTGTCTCCAATAAAAGGTTG<br>AATGTCGTCCTCAGTTTTCTGTCTAACGT<br>ATTCAGTAGTGGGTTCTGCCCTATTATTA<br>GCTATGACAACCCTAGAGTCTTCATCTAA<br>AATGCCGTATAGTAATTCCTCTTTTTCTG<br>AGGGCTTTCCTATGGATTTTCTTACTTTT<br>ACTGTAC |
| B-3 | 18528 | 18700 | + | intragenic | SSO_RS00095 (SSO0018) | Ferredoxins (Oxidoreductase activity) | AATCTCAGCTGTCGGTAATACGATTTAGA<br>AAGGATGAAATCGCTTATTCAATTTACTC<br>TAATAAGCGACTAGAGTCTATAAGAAACA<br>CTCTAGAGTATTCTACCATAGAGGGAGAA<br>GGCTTATTCAATGGATGCATTCTATGCGG<br>TAAATGTGTTAGTGTTTGCCCTTACGGT |
| B-4 | 20475 | 20647 | + | intragenic | SSO_RS00105 (SSO0020) | Archaea conserved hypothetical- Predicted nucleotidyltransferase | TTTAAATGTATATCCTCGCCCTTCAAAAA<br>TGCCTTCTAAAATCCTTTTACTTTTTAAA<br>ATCTTAAATGAAGAATAGTCTAGAGTCTC<br>TATTTCAGACTCTTCGACAGTGGATAACG<br>GTTTAGTGATCCCCTTTTTTCTTAACTCC<br>TCTAAGACGTAATAATGTTCTGGTTTGA |
| B-5 | 22703 | 22875 | + | intragenic | SSO_RS00120 (SSO0023) | Archaea conserved hypothetical | GATTTTTATGAATTAGGATTCACAAATAT<br>TAACCACTCCAGTCTACAATTAATATTAT<br>CTGCTTACTTCAAGAGTCTAGGCTATAGA<br>GTATTCGTGGAGTATGAGAAGAAAGGAAA<br>ACTACTTGATCTCTACGTTAGTGATGAAG<br>ATATGGGAATAGAAGTAGAATATGGTTA |

| | | | | | | | |
|---|---|---|---|---|---|---|---|
| B-6 | 24311 | 24483 | + | intragenic | SSO_RS00130 (SSO0025) | Archaea conserved hypothetical | GCTCGTAATTGAGGCCAAAATTCACAAAA<br>ATGATTACCTACAGACTCTAGAGTACTCT<br>AAATATTTCAAGTACGGTATGGCAGTATT<br>TCCATTTACTGGAGAGTGTAGAGTACCTA<br>AAGGTTGGATTTGTATATTTAACACCACA<br>AAGGATCAGTCTAGATTCTACTCTCTTC |
| B-7 | 26874 | 27062 | + | intragenic | SSO_RS00145 (SSO0029) | Archaea conserved hypothetical (cystathionine-beta-synthase CBS domain) | ATTTCTAAGATGAAAGAAAACAAGATGTG<br>GACTGTACCCGTTATCAAGGATAGGAAAT<br>TAATAGGTTTGATCTCTTATAAAGATCTT<br>CTTTCTAGAAGGGTGAGTCTAGAGACTAA<br>AGCGATAAACATTATGAGTCCCAGTGTTA<br>CTGTACAAATTGATGAGGATATTAATAGA<br>TTAATTGCAAAATTC |
| B-8 | 29213 | 29385 | + | intragenic | SSO_RS00155 (SSO0032) | Archaea conserved hypothetical | ATCGTCTTCCTTAATGCCTAAAGTTTTTG<br>ACGCATTCTTTACAAATAGGGTTCTATCA<br>AATTTGTAATCTTCGTATATATTCTTGAC<br>TTTAGACTCCAGACTCTTCACTACGTTAG<br>AAATTATCTCATTTTCAAAATATGCATGA<br>ATAGCAAAAACTAAACTTAACTTGCTTG |
| B-9 | 29661 | 29833 | + | intragenic | SSO_RS00160 (SSO0033) (*segC*) | *sso0033*<br>*segC* | ATCTTATTAACGTCCCAAATTTTCTCTAT<br>GATTTCCATACTTTTTTTCTTAAAATTTT<br>TACTTACGAATATTCTACTCTCTAGACCC<br>TTAGACTCTAGTAAGTCTTCTGGATCATA<br>ACCTTTGTTACTAATTCTTAAAATTATTT<br>TACCTCCATCACAATTAATACAAACTCC |
| B-10 | 30052 | 30258 | + | overlap end/overlap start | SSO_RS00160 (SSO0033)/SSO_RS00165 (SSO0034) | *sso0033/segA* | CAGAATAAATTATTGCACTATAGAGCTTC<br>TCAAACATACTTATACTTTTTAAAAATCT<br>AGCATATAAATATTATCATTATAATATGG<br>TGGACTCGTCAGTATTATCATGTAGAATT<br>CCTTATTAAACGTTATAGATAGAAGAGTC<br>TAGACATGATAGTCACAGTAATAAATCAG<br>AAAGGAGGAGTAGGCAAAACAACGACTTC<br>AGTA |

| | | | | | | | |
|---|---|---|---|---|---|---|---|
| B-11 | 30551 | 30723 | + | intragenic | SSO_RS00165 (SSO0034) | *segA* | CAATGTTGGTTGCTGACAGAATAGTTTCA<br>CCGGTAACACCACAACCCTTAGCACTAGA<br>GGCAATAAAGAATCTCGACTCTAGATTAA<br>AGAGTATAGGGAAGAACGCTTATTCTTTT<br>ACAAATTTTTCAAAAAAGGTAGTTAAGCT<br>AGATAATCTATCATCAGTAAAATTCACA |
| B-12 | 30730 | 30909 | + | overlap end/overlap start | SSO_RS00165 (SSO0034)/SSO_RS00170 (SSO0035) | *segA/segB* | ACAATACCACCCTCTAGATTATTCATTGA<br>AGCTTCTAGACTAGGAGTTCCAGCGTTAA<br>GATATGAGGAAGTTAGAATAAAGAAACCT<br>AAGCTAGCTAACTATTATCAGCAATTAGC<br>AAAGGTGATTAGTGAATGAGTGAATTAGA<br>TTTCCTATTGAAGAAGAAAAGAAAAAGTG<br>AGGACG |
| B-13 | 31166 | 31346 | + | overlap end/overlap start | SSO_RS00170 (SSO0035)/SSO_RS00175 (SSO0036) | *segB*/archaea hypothetical | TGCAAAAGAGAGATTTAAAGAGTAATATC<br>TGGTAATATTTCTAGGAAGTATTACTAAT<br>CAGGTTTAAATGCTAGAGATTATATTTAT<br>CTGTAACTGTTAAGGTGATTAGAATGGAC<br>ATATGTGTAAAGGCTAGAAACGATGAAGA<br>AGCTACAAGGGCTCTACAATTAAATTACA<br>ATTGTGT |
| B-14 | 31693 | 31865 | + | overlap end/intergenic | SSO_RS00175 (SSO0036)/SSO_RS00180(SSO0037) | Hypothetical protein/ flagellar hook-basal body protein | AATCTCAAAAAATTAGGATTATATAAAAA<br>TAGGATTATATAAAAGGGAGATTTTAAGT<br>CTAGAGAGTGTCACTCTTGACTGAAGAAA<br>GACGAGTATAGAGCGAGGAGTTAGCGGAC<br>TTTAAAACCGCGTTGAGTTACAAAAATAT<br>ATGCATAGATAGCGTTTAGGCATTGAGG |
| B-15 | 34685 | 34857 | + | overlap end/overlap start | SSO_RS00185 (SSO0038)/SSO_RS00190 (SSO0039) | Conserved PadR family DNA-binding transcriptional regulator /conserved Phosphomethylpyrimidine kinase (Thiamine biosynthesis) | TTAAAGAATGTCTTGTGAAATATATTTGA<br>GTAATTCTAGAGATTTATTTACAACAGTG<br>ACTGGATCTCGATCTAGAATGAAGAGTGA<br>AGGCTCTTTACCATAATCACCTAAATCCA<br>CAATATAATTAGGAACCTTAGACAACTTA<br>CGAATACACGAATCAACCATAAAGTTCA |

| | | | | | | | |
|---|---|---|---|---|---|---|---|
| B-16 | 43657 | 43829 | + | intragenic | SSO_RS00245 (SSO0049) | Conserved PadR family DNA-binding transcriptional regulator | AAGAGGGTAGAAAAGCTATAGGAGCTATG<br>TCAGAGGAGGATAAAATCAAAGAAGCAAT<br>AGAACAATTGGAGTTCTCCGCTAGATACA<br>TAGTCGAGAATCTAGAGAAATTAAATGAC<br>GAAGATAAGCGAAAAGTGAAAAATATATT<br>GGACGAATTAAGTAAAGTTATGCGATAG |
| C | 120356 | 120528 | + | intragenic | between SSO_RS00685 & SSO_RS00690 | CopG family transcriptional regulator/hypothetical (could be socitrate dehydrogenase (IDH)) | TTATATGTCACATCTAATATTCCCCCTAG<br>TCTAACAATTTTACTCTTCATTCTGATAA<br>AAATTCAAGTAAGAAGAGTGAAGAGAGAA<br>GAGAGAAGAAAATAGAAGGGAAGAAGAAG<br>AATAAACGCGGAAGAGCGCAAAAGAGAGC<br>GTTTGAACCCACGGGTGGGCCCAATAGC |
| D | 1001960 | 1002132 | + | intergenic | between SSO_RS05720 (SSO1161) & SSO_RS05725 (SSO1162) | Conserved hypothetical containing HEPN domain/MFS transporter (carbohydrate transport) | CTTAAAACAAGAGAAAAGTAGAGAAATAT<br>TCTATTATACTCTCTTTTTCCTAAATTTT<br>AATCTTAAACTTAAGGGAAGTAGAGTCTA<br>GACTCACGTTAGGGTATGAAATTATCTCA<br>TCCTAATTTTTATTTGGCATAACGCTAAT<br>ATGAAAAACATAAATAGTTAGAGTGTTA |
| E | 2055588 | 2055760 | + | intragenic | SSO_RS10935 (SSO2241) (*bps2*) (*clsN*) | BPS2 protein homolog - AAA family ATPase (SMC like protein) (ClsN) highly conserved in archaea and in bacteria | TCGTCTATTTGGAACTTAACAGAACCTAT<br>CCTATTAATTAACTCTTCCTTTTTAGCCT<br>TTAATTTTGCAATTTCATTCTCTAGAGTC<br>TGTAACTCCTTTAATCTAAGGTCGATTTC<br>TTCTTTCCTTTTCTGTAATCCTATTATAT<br>CATTACGCAAACTGTCCAAGAGCGAGGT |
| i | 50316 | 50488 | + | intragenic | SSO_RS00305 (SSO0060) (*gnptA*) | Glycosyl transferase (GlcNAc-1-P transferase) | TACTTGATACTCCTTAAATCGACCCATTT<br>TCATTACTATATGTGTTAATGAATGAGGT<br>TTTTCATCCCAATATAATCTACCGCTAGA<br>GTCTACTTTGCCAAATGATACTCCCTTGA<br>AGTTAGTTCTTAGCTTCAATATAAATTCC<br>ACGACGTATGGTATGTAGAGAATTGCTA |

| | | | | | | | |
|---|---|---|---|---|---|---|---|
| ii | 109363 | 109535 | + | intragenic | SSO_RS00625 (SSO0129) (*cbiG*) | Protein G (CbiG) (Cobalamin biosynthesis) | TACTATATAATCTATTCTATCGTTTAGAA<br>TATAAAGTGATTTTAAGGTCATCCTTAAG<br>GAATAAACAAGAACATCTAGGGG<u>AGTAGA</u><br><u>TTCTAGA</u>TACCTTATACCAATGTAAATTC<br>CAGAAGGTATTAAGTATAGTCTATTACTG<br>CTCTTATCTTCCATACTATCAACATTTT |
| iii | 196087 | 196259 | + | intragenic | SSO_RS01130 (*rpoB2*) | DNA-directed RNA polymerase, subunit B" | ACTATAACTCTTTCAGATCCATTTACTAT<br>GAAATACCCTCCTGGGTCTTTAGGATCTT<br>CACCTATCTCAATTAGCTTA<u>TCTAGAGTA</u><br><u>TACT</u>GTGATATTGGGTCTATGGCTGATTT<br>AAGCATTATAGGCAGGTCGCCTATATAAA<br>CCTCTTCTGGTTCTGCTTCAATATTATT |
| iv | 235204 | 235376 | + | intragenic | SSO_RS01360 (SSO0274) (*tgtA*) | Queuine/archaeosine-tRNA-ribosyltransferase (Glycosyl transferase) | AGGTAAATCTAATATTACTCCGATATCTG<br>GTTTTATTTTAAGCTGATAATTCACGATC<br>TGGAGATTGGTTATTCCTATTTC<u>TCCATA</u><br><u>CTCTAGA</u>ATTTGATATGCCCCTGAATCTG<br>TCATTATGATCATTTCTTCAGAGCGTAAC<br>TCCTTATGGATATCATCCTTTATATAGT |
| v | 254482 | 254654 | + | intragenic | SSO_RS01490 (SSO0298) (*thiF*) | Thiamine biosynthesis related protein | GAAAAGGATACTCATTTAGAACGCATGTC<br>TTACATTGTGAGTTCTTCTCGATGTTAAT<br>TCTCTCAATTTTTAATTC<u>TCTAGAATCTA</u><br><u>TA</u>TAGAATAATGAATAATCCGGATTGCCT<br>CTCAAGTGATTAAGCATTAAGTTAACTTG<br>AAGTGTAGCTGTTAATTCTACTATTAGT |
| vi | 717556 | 717728 | + | intragenic | SSO_RS04240 (SSO0873) | Polysaccharide biosynthesis | ATTTTGGGAGGAGTACTCTCAGTCCTCCT<br>TAACTACTCTTTTCTGTCTAGAAAAGTGG<br>GGCTAGTCTTACCTTC<u>TCTAGACTTTGCA</u><br>TTTCTGTTTAAGCACTTCAAGGAAGGCTT<br>ACCTCTCTACTTGTCTTCTTCAGCTAATT<br>TCCTCTCATCGCAAGGGGACAGAGTTAC |

| | | | | | | | |
|---|---|---|---|---|---|---|---|
| vii | 818763 | 818935 | + | intragenic | SSO_RS04795 (SSO0963) *topR-2* | Reverse gyrase | AAAATTGTTTCCTAACACTACCTTTACTT<br>CATTATCTAATACAAATTTATCTGAAATA<br>AAAATTAAACTGGAAGACTCTAGAAAAGA<br>AGAAGGGAATAGGGTCCATTTTAACATAT<br>CAACTGGTTTACTTATAGTAGAGTCTCCA<br>ACAAAAGCAAAGACTATAGCTAAAATGT |
| viii | 965294 | 965466 | + | intragenic | SSO_RS05535 (SSO1119) | Highly conserved protein of unknown function DUF1641 | TCATAGTCTATCTGATCAATTACTTTCAA<br>CACTTTTTCGTCAGTTAGCTTCTTAATTA<br>TTGGCGATAATTTCTCTAACGCGGTAAGA<br>GTAGAGTCTATATCAATCTGCGATAGTAA<br>TTGCAAAGCCCTTTCGCTTGTTAGTTTTT<br>CTAGTATTGGCCATATTGCTTGTATTTT |
| ix | 1066884 | 1067056 | + | intragenic | SSO_RS06045 (SSO1228) (*tmoA*) | Toluene-4-monooxygenase system protein A (containing YHS domain) -toluene catabolic process (Ferredoxin) | ATATTTGGTAGCGAAAGGATTCTGGCTCT<br>CCGTTAGGCTAATAGGAGCCTTAACCGGA<br>GTTTCAATGGATTACCTCACTCCTCTAGA<br>CGCTAGAATAATGTCATACAAGGAGTTTA<br>TGACAGAATGGGTAGGTGCGCAGTTAAAG<br>AGATTATTAGAAGATTATGGAATAAGAT |
| x | 1297295 | 1297467 | + | intergenic | between SSO_RS07065 (SSO1444) & SSO_RS07070 (SSO1445) | CRISPR-associated protein Csa3 (Type I-A) /CRISPR-associated transcriptional regulator Csa3 | TATAAGAGATTATCTCAGTCTTATTGTCG<br>AGGAGTAGTGGTAGGAGTGATACCTTTCA<br>ATTCTATAAGAGATTATCGGAAGAGACAA<br>GACTCTACAGTACAATATGCCCCAGATGC<br>TTTCAATTCTATAAGAGATTATCAACCGG<br>AAGTACGAACTATGTTGCTGAGATTGTG |
| xi | 1699509 | 1699681 | + | intergenic | between SSO_RS09090 (SSO1883) & RS_SSO09095 (SSO01884) | Transposase ISC1234/archaea conserved hypothetical (DNA polymerase beta domain protein region) | GCCAATAAAAATTCAATTTTTTACATTCC<br>GGACACTCTCACGAGTTAGTCCACAATTA<br>CAGACTAATAGATATTGAAACTGCTTATA<br>TAGATTCTAGATACGGTAATTTAGAATAT<br>GAAAGTGAAGACTTGAAATCACTAATTCA<br>AGTTGCAGAGAATGTAATAAAATCACTAG<br>AGG |

| | | | | | | | |
|---|---|---|---|---|---|---|---|
| xii | 1910560 | 1910732 | + | intragenic | SSO_RS10195 (SSO2094) (*treX*) | Glycogen debranching enzyme - glycogen catabolic process & protein homotetramerization | TAGGGCACATCCAGCATTCAGAAGGGAAA<br>GATATTTTCAAGGAAAGAAATTATTCGGC<br>ATGCCGTTAAAAGATGTGACCTTCTATAC<br>TCTAGAAGGTAGGGAAGTTGATGAGAAAA<br>CATGGAGTTCCCCGACGCAACTAGTTATT<br>TTCGTGTTGGAGGGAAGTGTTATGGACG |
| xiii | 1999893 | 2000065 | + | intragenic | SSO_RS10630 (SSO2176) | Hypothetical (present in some crenarchaeota only) | AACGGTAAATAACTATATATTGTTGGGGA<br>TGGTGATTCATTTTCAGAATTTATCTGGA<br>ATATATATATATGTACGTTTTTACTAGTA<br>CCGTTTGCTTGATAAAGTATATTGTTGTG<br>ATAAATTAGGTTTAATACTATTTCACTCC<br>TATTTAATTTTAAAAATCCCTTATATAC |
| xiv | 2112188 | 2112360 | + | intragenic | SSO_RS11245 (SSO2309) | Cofactor biosynthesis protein (heme biosynthetic process) | GATTGGGCTAGCGTTGTAAACATAACTCG<br>CCATGTTATCTATTATTTTCTCTTTCACC<br>ATTTTTCCATTCGACCAACTGAATAGTGA<br>AGGCTCTAGATAACAAGCAAGAGCACATG<br>CATTACAGCTCTCATATTTTTCCCATGGA<br>TAACTGTTCCATAATTTTACTATATCGA |
| xv | 2281328 | 2281500 | + | intragenic | SSO_RS12200 (SSO2514) | 3-hydroxyacyl-CoA dehydrogenase/enoyl CoA hydratase (Lipid metabolism) | TGGGGTTTTCTTTTTTCTAAGAAAGCTTT<br>CACTCCTTCTTCTACATCTTTAGTTGTGA<br>ATAAAAGCCCAAATAATGTTGACTCTAGA<br>GTTTGTCCAGTCCAGATATTAGATTCGTA<br>TCCTAACTCTATTGCTAATTTAGCTGCCA<br>ATAGTGATATTGGTGATTTTTCAGCTAT |
| xvi | 2377298 | 2377470 | + | Overlap start/overla p end | SSO_RS12665 (SSO2610)/ SSO_RS12670(SS O2611) | Conserved hypothetical/conserved hypothetical in archaea & bacteria | TACTAGTTCTCTCAAAGATATGGCTTTAA<br>GTTTAGAATAGAGCGGTGGATAGTAAGCA<br>GCGTAAAAGATAAAAGGTATTCCTTTACC<br>TTTAGAGTCTAGATCCATCATTAAATATA<br>ATTACGTATCAAAAGCATATTAACTTATC<br>CACAACATAATATTGTAATTTAACTAAT |

| | | | | | | | |
|---|---|---|---|---|---|---|---|
| xvii | 2589275 | 2589447 | + | intragenic | SSO_RS13735 (SSO2828) | Hypothetical conserved in crenarchaeota | AATAATGTGGGTAATATTATAAGATCCTA<br>CATATTCTCTACCTTGGAAAACAACCGGT<br>AATTTCCCAATATCGCCTTCACATCTATA<br>GAGTCTACTATCAGAGTTTAGGAGAAATT<br>TTGTAACTGGAAGAACTAAAGGGATGAAA<br>AATTCTGGATCGCTTAATATTTTAATTA |
| xviii | 2630263 | 2630435 | + | intragenic | SSO_RS13955 (SSO2877) (*acd-6*) | Acyl-CoA dehydrogenase (lipid metabolism) | CTTTTTCTTCTGTTCTTCATTACCGAAAA<br>GCAATATTGGAGTCATGAATAACCCTCCA<br>ACAGATATTCTCGTAGAAAGTGAGGCCCA<br>GACTCTAGATATCTCTTCTTGAGCTATGG<br>CAGTCATTAGCGTATCTCCACCTTGACCA<br>CCGTATTGTTCTGGTACTGCAACACCAT |
| xix | 2727724 | 2727903 | + | intragenic | SSO_RS14490 (SSO2984) | Hypothetical conserved in archaea & bacteria (nickel permease) | TTGATATAGGGGCAAACAAGGCTGGGTTA<br>TTACAAGGAGCTTTCGTCATAACTCCAAC<br>CAAGGTGTTGGTTATAGTTATCGCTTCCA<br>CTGCTTATAGTATACTCTACTCTATAGAG<br>GTAATTTCAGTATTTATAATAGCATCTGC<br>AGTTTCAATTATATCTCTCTCACTATTAA<br>ATTTCG |
| xx | 2967544 | 2967706 | + | intragenic | SSO_RS15595 (SSO3218) | ATPase | AAAACCATCGTGATCGACATAAACGTTTT<br>TTGCAATAGATCTCAATGCTTCTGCCATT<br>ATGTTGTAATCATATCTAGAATCTACCCC<br>AGCCAAACCTACATAAGCTACATCTGGTT<br>TCATACCTTTTGTGGCTAATAAAATCGCT<br>CTGTTAACGTTTTTTACA |

**Table S1. SegB binding sites identified by ChIP-seq and DNase I footprint on *S. solfataricus* chromosome.** ChIP-seq enrichment peak sequences and relative genomic coordinates, with high-enrichment peaks labelled from B to E and low-enrichment peaks designated by Roman numerals (i to xx). Sites determined within the peak sequence through MEME are underlined and highlighted in yellow, whereas those further identified by DNase I footprint are underlined and highlighted in blue.
