## Supplementary Table S2 for "Coupling chromosome organization to genome segregation in Archaea"

| #Peak ID-Figure 6 | Start | End | Strand | Position of the peak | Gene accession number/name | Product (gene function) | DNA sequence under peak Motifs identified by MEME (yellow) DNaseI footprint (blue) |
|---|---|---|---|---|---|---|---|
| 1 | 62063 | 62362 | + | intragenic | SACI_RS00365 (Saci_0079) | Aminoimidazole ribonucleotide synthase related protein | CGTTACAAATGGCGGTATAAGGGCTACTG<br>CAAATGAGTTGCCTGAAAACTTATCATTA<br>GCCATAAACACAGAATCTTTTCTGAGGCT<br>GATAAACAGAAAAGTGTTAAATATGCTTA<br>ATGAACTGCAGATAGATGTTTTCGGGCTA<br>TCTATAGACTCTATACTCATCTTCACTGA<br>TATAGGTAATGAAGTTGTAAGTAAATTAA<br>AAAGCAAAGGTATTAATGCGGAGATAATT<br>GGAAAAGTAGTTCAAAGACAAGATTCGCC<br>ATTAATTACTAGTGAAGGTAAACCTCTAA<br>TCATGAATTT |
| 2 | 153097 | 153396 | + | intragenic | SACI_RS00905 (Saci_0188) (*aroA*) | 3-phosphoshikimate 1-carboxyvinyltransferase | AGAGGAGATGTTCCAGGAGACTATGCATT<br>AGCTTCATTTTATGCTATTGCCTCTGCAA<br>TAACTGGAGGAGAGATTCAAATTAAGGGA<br>CTCTATTCTCCTCCATATTACGATGGCGA<br>TCACAGCATTGTAAAAATAATTAAGAATA<br>TGGGTGTAGACTCTAGAGTCGAGGGTAAC<br>TCATGGATTGTTTCAGGGAACGGAATTAT<br>AAAGGGAATAAAAGTGGATGTGGACGATA<br>TGCCTGACCTTGCACCATCTATAGCTTGT<br>ATTGCACCCTTTGCTACCTCAGAGACTGA<br>AATTACTGGA |
| 3 | 154065 | 154364 | + | intragenic | SACI_RS00910 (Saci_0189) (*aroD*) SACI_RS(Saci_0190) | Type I 3-dehydroquinate dehydratase | AAGCATATCTTAGAAAAATAGCTGTAATA<br>GCTAAGAGAGGTTATAGGGAACTTTTGAT<br>GAGAGTTCTTGATAATTACGATAATGCAG<br>TTGTTATGCCTATGGGTGTTAATGGAGTA<br>GAGAGAATAGCCTTCTCTCTTCTGGGATC<br>TAAGTTGATATATGCCCATGCTGGCGAAG<br>AAACTGCTAAAGGGCAGTTACATTATAAG<br>GATGTGAGAAGAATCTTAAATCAACTTTC<br>CACAATTATGTCTTCACCATCAACCTGAA<br>CGCGGTATAGTTTCAATGGTTTCTTGGTT<br>GCTCCTTTGA |

| | | | | | | | |
|---|---|---|---|---|---|---|---|
| 4 | 156185 | 156484 | + | intragenic | SACI_RS00925 (Saci_0192) | ATP-dependent DNA helicase | CTGACTATCTTCTAAAAATATACTTTCAGGCTAAGGCAAATGTTTTGGTAGTATTTCCGTCATACGAGATAATGGATAGGGTTATGTCTAGGATTTCATTACCTAAGTATGTTGAAAGTGAAGACTCTTCAGTAGAGGATCTATACTCTGCAATATCTGCAAATAATAAAGTCCTGATAGGGAGTGTAGGAAAAGGAAAATTAGCCGAAGGCATAGAATTAAGGAACAATGATAGAAGTTTGATTTCTGATGTGGTAATTGTGGGTATACCTTACCCACCACCTGATGATTACTTGAAAA |
| 5 | 158493 | 158792 | + | intragenic | SACI_RS00945 (Saci_0196) (*mpg1*) | NDP-sugar synthase | TTCCTAAATTATATAATGGTATCTGCTATAGTTCTTGCAGGTGGATACGCAACTAGACTAAGACCATTAAGCTTAACAAAACCTAAAGCACTTCTTCCTGTACTAGGAAAACCATTAATGGACTATACACTCTACTCTCTAGCATCTTCAGACGTTGATACGATATATCTCTCCTTGAGGGTCATGGCTGATAAAGTTCTCGACCATGTTAAGCAGTTAAACTTACAGAAAAACATAGTTTCCGTTATAGAGGAGAGTAGGCTAGGAGATGCTGGTCCACTCAAATTCATAAATTCAAAG |
| 6 | 159603 | 159902 | + | intergenic | SACI_RS00945 (Saci_0196) (*mpg1*) / SACI_RS00950 (Saci_0197) | NDP-sugar synthase/ conserved hypothetical protein | TTAATCTCAGTAGAATTTGTCAATGAAATTGCCAGCATTACTCCCTATTTCTCTTTTTATAACTATCCTAATCCCACTCTGCATTTTTTCGATTGTCACAGAGTCTTCAGTACATACTCTAGAGATCCTAATAAACGGTCCCTTACAGGTAACTATAACGCAGGGAAATAATTTCACTGTAATTAGCAAAACTACCTCATTGAGATTACAGGGTAATATAACTATTCAAGCATCAACCCTTCAACCTGGATACCAGATATTCATTAATGGCATCAAGGCAAATACGCTAACCCTCAATTC |

| | | | | | | | |
|---|---|---|---|---|---|---|---|
| 7 | 160700 | 160999 | + | intragenic | SACI_RS00955 (Saci_0198) | Thioredoxin domain-containing protein | TAGGGTTGGTTTCAAGAGACTTCTTAATG<br>AGATTATAAGATTATGGAAGAATGAGAGG<br>GATAAAATATTTCAGACTGCTAATTCTCT<br>TCACTCTTCACTTCAGAATCTAGGATATC<br>AAAAAGTTGAAGCCCAATGGGATCAAGTG<br>GAGAGCATAGTGAGTTATATTGCATCAAA<br>TTTTGACTTTCAAAACGGTGGACTCTTAG<br>GTAGTATGAAATTTCCTCACCCAACTATA<br>GATCAGCTCTTAATTGCTTACTCTTTCTA<br>CACAAAGAGTGATACTGAGGCTAAACTTT<br>CTATGTTTAC |
| 8 | 161351 | 161650 | + | intragenic | SACI_RS00955 (Saci_0198) | Thioredoxin domain-containing protein | CAATGTTGATACTGGAAAAGAAGTTGAAG<br>GCAGAAAAGTGTTAAGAAGGAATTATGAT<br>TTAAGAGAATTAAGTAAAAGGTTCAAGGA<br>TCCAATAGGTAAACTGAATGATGTTAGGG<br>AAAGATTAAGAGTCTATAGAGAAGAGAGA<br>AGAAAGTATCCCTTTATCGACACAAATGT<br>ATATACTCACTCTAACTGTAGAACAGCTG<br>AGGCTTTAACATTAGCTTATCCAATTACA<br>GGTAAAGGTTTAAATGAAGCCCTTAAGGT<br>AATAGATATGATAAACACTAAGATTACTA<br>GAAGACTAAC |
| 9 | 161892 | 162191 | + | intragenic | SACI_RS00955 (Saci_0198) | Thioredoxin domain-containing protein | TATTCAATAGATTCCCCTTACATTGCAGG<br>TGTAACATTTAACTCAATGGCTTTTGAGA<br>AAGGTTTAGCTCATATTGTGGTAGTTGAC<br>GAAAAAGACGGTAAGGCTAAAGATCTTCA<br>CCTATCTGCACTGAAAATATATCATCCTT<br>TCAAGGTCGTGGAGCTCGTGAGTGAAGAC<br>TCTATAGATATGCTTCCCTCTTTTATAAA<br>ATCCATGGTTAATTACAATAAAGGTTCCA<br>GTAGAGTTTATGTGTGTATAGGTAATACT<br>TGTAATTTACCGGTAGACTCTTCAGAGAA<br>GATTAGATTA |

| | | | | | | | |
|---|---|---|---|---|---|---|---|
| 10 | 162780 | 163079 | + | overlap end/ intergenic region/overlap start | SACI_RS00960 (Saci_0199) / SACI_RS00965 (Saci_0200) (*nucS*) | Short-chain dehydrogenase/ SDR family NAD(P)-dependent oxidoreductase/ endonuclease NucS | GAAAATTAGGTGATGCAGGTGCACCGCCT<br>GAAGATTTTGCTCAAGTTATAACATGGTT<br>ATTGAGTAATGAGGCACAATGGGTTAATG<br>GAGTAGTTATTCCAGTTGATGGCGGTGCT<br>AGACTGAAGTGGGTGTAGAGTACATTCTT<br>CTCTCTAGAGTGAATATATATTAGTTTTT<br>TATAAGTAATACTATGTTCAAGGTATTAC<br>TCGAGCCTGATTTACAGGAGGCTTTAATT<br>TTCCTCAATGAATCGGTGAATGCTCTACT<br>TACAATTTATTCTGAATGCGAAATCTTAT<br>ATTCAGGCAG |
| 11 | 163122 | 163421 | + | intragenic | SACI_RS00965 (Saci_0200) (*nucS*) | Endonuclease NucS | CAAACCAGATGGGAGTGTTATAATCCACG<br>GACCTACTAAAAGAGAGCCTGTAAATTGG<br>CAACCTCCAGGATCGAGAATAGAGTACAG<br>TATAGAGAGTGGAGTATTGACAGTAAATG<br>CTGAAAGGAAGAGACCAAAGGAAAGACTC<br>TCCATTCTGCACCACAGAGTCTACTATAT<br>TACCTCTTCAGAGGTTAAGCCTGGAGAGT<br>TCTTTCTAGTGGGAAGAGAGAAGGATGAA<br>GTGGACTTCATTATAAATAACCCTGATGT<br>AATAGAGGGTGGATTTAAGCCAATTCACC<br>GAGAGTATCG |
| 12 | 165363 | 165662 | - | overlap end/overlap start | SACI_RS00980 (Saci_0203) / SACI_RS00985 (Saci_0204) | *segA/segB* | TGAGTTTGGACTACTCTTCACTTCCTCTA<br>AATCTTCCCTAGGGCTTTTTGAAGCGTCA<br>TTCTCATTAACCTTTTCCTGGGTTTCACT<br>AGTTTTGGTCTCTGCTAATTTCTGGGTAG<br>TAGACTTGGAAGTCTCCTGTCTGGACTCA<br>ACTTTCTGTTCACTAGCCTCTACTTTCTG<br>TTCAGTCTTTTTTCTGTTTAAGATTAAGT<br>CTAACTCACTCATCCTTCTATCACCACTT<br>TAGCTAAGTCTTCATATAATTGTGAAAAC<br>TTAGGTCTCTTCACTCTAAATTCTTCATA<br>TCTAAGGGCT |

| | | | | | | | |
|---|---|---|---|---|---|---|---|
| 13 | 165663 | 165962 | - | intragenic | SACI_RS00985 (Saci_0204) | *segA* | GGAACTCCGAGTCTAGTGGCTTCAGAGAA<br>AAGTTTAGACTGTGGGATTGATAAGTCTA<br>TACTCTTGACTGAAGGTAGCTCAAGCTTT<br>ACAGATTTCTTACTCATGTTCGTAAATGC<br>TATTGCAGGTTTCCTTAGACCCTGCAGTC<br>TAGAATCCAGATTCTTGGCAGCCTCTAGC<br>ACAAGGGGTTGTGGGGTAATTGGTGTAAC<br>TATCTTATCCCCAGCAATCATGGCAGAGA<br>CTGCTAATGTGCCCAAGTTAGGGGGAGTA<br>TCTATGACTAGAAAGTCAAAGTTCTCGGC<br>AAGTTTCTTA |
| 14 | 165961 | 166160 | + | intragenic | | *segA* | TAAGAGAGGAGACTATAGACTCTATATCA<br>CCATTGAGCTCTAGTTTGAGTAGACCAAT<br>ATGTGCAAGAAATACTTCCACGTTAAATA<br>TATTCACACTTTTTCCTCCTAATGGATAT<br>TCCCTTTTTTCCCTCTTTATTCCAAATGA<br>TATTGTAGCTCCACCTTCGGGATCAAGGT<br>CTAGTAATGCCGTATTTTTATTTTTG |
| 15 | 166138 | 166437 | - | overlap end/ intergenic region/overl ap start | SACI_RS00985 (Saci_0204) / SACI_RS00990 (Saci_0205) | *segB/*(Fe-S)-binding protein (oxidoreductase) | GTAATGCCGTATTTTTATTTTTGGATAAA<br>GTGTAAGAGAGATTTACGGAGGTTGTAGT<br>TTTCCCTACACCTCCTTTCTGATTGATGA<br>CAGTGATTATCATATATAGACTCTTAACT<br>CTATACCCTTTATTTCAGATTCTTCTCAT<br>AATACACCGTAGTCTGGAGTGAAAAATCT<br>AAAGTAAATCCACTAGGTGAATAGGTTTT<br>AAACCCATTTTTATAAAGGCGAGTGTGCA<br>TGACGGATTTGTTGAAATCACAGTATTAA<br>CCCCACTCTTCATTACTTTCTCCTTCTTT<br>CTACTCATTA |
| 16 | 166775 | 167074 | - | intragenic | SACI_RS00990 (Saci_0205) | (Fe-S)-binding protein (oxidoreductase) | GTTCTCACCTACATGTGAGTGAGCTAAAC<br>CACAGCAAGTATTAATGACTTTTACGTTG<br>TACCCTAGACCTTTGAAGTATTTCACTGC<br>TTTCTCTACACTACTTCTAGATATAACTG<br>AGGTTAAACATCCTGGAAATATTATTAAA<br>TCTTCATTCTGGTCTCTGTACTCTAGAGG<br>AAGGCTGGTCTCCACTCCCATTCCTATTT<br>TCAGAGGTAAGGCTAATGATGAGGGTTTT<br>TCAAGTAATCTTAACATAGATCTTTCCAA |

| | | | | | | | |
|---|---|---|---|---|---|---|---|
| | | | | | | | TGGGTTAGACTTTCTTGAATTTACGAATA<br>TCTCTGAGTA |
| 17 | 167624 | 167923 | - | intragenic | SACI_RS00995 (Saci_0206) | FAD-linked oxidase C-terminal domain-containing protein (glycolate oxidase) | ATCTTCGCTAAAACTTCAGGTAAGTTACT<br>TCTCAACACATTACAGTCTAAGGTGATAT<br>ATGCCGGTGAGATAACTCCCATAGCCGGA<br>AAAGCTCCCCTTCTTGCACTCCAGAACTT<br>TTCTGGTTCCTTTGGATTAATTATCTCAC<br>CGTTATTTTCGCTTATTACATCCCTGACT<br>CTACTCTCTTCACTCTTCACTTGAATCTC<br>TGAACCATCTAGTTCTATAAGCAATATCG<br>CCTCAACTTCAGGTAAGCCTGCCCTATAT<br>CTACTCTTTTCTATTGCAATTATGGAATA<br>TCTGTCCATC |
| 18 | 168425 | 168722 | - | intragenic | SACI_RS00995 (Saci_0206) | FAD-linked oxidase C-terminal domain-containing protein (glycolate oxidase) | CCGCTGAGACTAGTTCCAGAACCTCTAAT<br>TATAATTTTTCTTCTATTCTGTATCAAGT<br>ATCTTATTACAGCAATTGCCTCCTCTTCA<br>TTACCGGGTAGTACAACTAATTCTGGTTC<br>TCCCTTTACTGCTGTTAGACCATCAAAAC<br>CATAGAGTCTTCTCTCTTCATCCTTTATG<br>ACCCACTTGTCCCCAACTATTTGTTTTAG<br>GTCTTGGATCAACACTACAGTTTCACTTT<br>ATGATTTAATGGAAATAAATTTTTGTTAA<br>CAATTAATTAGTAGGATTAAATTATAGAA<br>AGAACTTT |
| 19 | 169202 | 169501 | + | intragenic | SACI_RS01000 (Saci_0207) | FAD-dependent oxidoreductase | ACCAAAATTCTCCAGTGGCTATAAGATTT<br>GGAAATCACTGTCAGGATCACTAGGATTA<br>TTAGGAGCCTATTTAGAGATCATTGTAAG<br>ACTGATACCTAAACCAGAGAAGATAATGT<br>ATGCTGAAATAAACGACGTAGATAAGGTA<br>TTAAATGAGAGACCGTGGGGAATACTCTT<br>CTCTGCAGACTCTGGAGAAATAAAGAAAT<br>ACGCTATATTCGCTGGGTTTCAAGATTAC<br>TTGAGGAGTGTAGAGAAGGAGTACAATAT |

| | | | | | | | |
|---|---|---|---|---|---|---|---|
| | | | | | | | TTCTCTTGTTGATGGCATTCCTAGCCATG<br>ATCTACAGTG |
| 20 | 174483 | 174782 | - | intragenic | SACI_RS01025 (Saci_0212) | Cystathionine beta-synthase domain-containing protein | ATGGAGTCTATTATTGCCCTTGAATTGCC<br>CTCCTTAAGTTCACCAGTTAACTTAGCCT<br>CTATCGGCATAGAAAGTTCGAATTTCCTT<br>GCAATAACTTTAAGCGCATCAATTCCGCT<br>AAACATTCCCACAATCATGTTACCTTCTA<br>GTACTGGCATACCTGATATCTTTCTGGTG<br>AGTAGAGTATTCACTGCAGAGTGAAGATT<br>CTCTGCTCCATTTACAGTTATTACGGGGT<br>AGTTCATTATTTCCTTTATCGGCATTGCC<br>ATTAATCTCTCTTCCTCAGTAAGTATGGA<br>GGACTTCTTC |
| 21 | 296995 | 297294 | + | intragenic | SACI_RS01710 (Saci_0350) (*gcs2*) | Glycine cleavage system protein GcvH | GGTGAAAATAGAGGGAAGTGATATTGCGG<br>TAGTTGGTATAACGGATTTAGCACAAACC<br>ATGGCAGGGAAAATTGTGAAGATAAGGAT<br>AAAGAAGAAAGGTATAAAAGTAGAGAGGG<br>GAAGACCTGTAGCAACTCTAGAGAGCGGG<br>AAATGGGCTGGACCAGTGCCTGCACCAGT<br>TTCCGGTGAGGTAGTGGATTCCAACTCAG<br>AAGTTGAAAAAAGCCCTGTGATCTTAAAC<br>AGAGATCCTTATGGTCAAGGATGGATCGC<br>AAAGATTAAAATTAGTAACCAGGAGGAAG<br>TTAAGCAATT |
| 22 | 404080 | 404379 | +/- | overlap end/overlap start | SACI_RS02330 (Saci_0482) / SACI_RS02335 (Saci_0483) | MarR family winged helix-turn-helix transcriptional regulator/ helix-turn-helix domain-containing protein | TGGGCCTGATAGAGGTCCAGGAAGAGAAG<br>ACGAGGGTAAAGAGCGTAAAGAAAATAGC<br>CAAACTAACAGACAAGGGCCGCATGGTCT<br>TTGAGAAGATTCTAGAGTTAAACCAGCTC<br>CTCTCAGAAGACAAAGCATGAATAGGCGT<br>CTTACGAGCTCGATAAAGACTTCGAGGCT<br>TTTTTCATTTCGACGATCCTTTCAATCTC<br>AACTAATAATTGAGCGACCTCTTTCCCGC<br>TCTCTGTCAATATTACGAACCTGCGGCGT<br>GGGAACGAGTGCTCTCTTTCTTCATTAAC<br>GAGGCCTAAC |

| | | | | | | | |
|---|---|---|---|---|---|---|---|
| 23 | 502140 | 502439 | + | intragenic | SACI_RS03005 (Saci_0632) | Conserved hypothetical protein | GAACATTATTTAGCTGAGGTCTACTTCTC<br>GTTCTGTCCACACTTTGAGCCTAACCACA<br>GGGCGGAAAGTGTAGTAGAATGCCAGTGG<br>CTCTTCTATGATATCGATAACGTGAACGA<br>AACGGTGCTCAGGAACCTCTCTAGACTCT<br>CGCCTAGACCAGTTGTGATCCTCTATTCG<br>GGGCATGGTGCACACTTATATTACAAGTT<br>GTCCCGGAAAGTACGCACAGAAGAATACA<br>AGAAGATCTGGCAAAGATTCGCTAAGCGT<br>TACTTGACACAATATCTAAATAAGATAGA<br>TAAAACGAAA |
| 24 | 580288 | 580587 | + | intragenic | SACI_RS03450 (Saci_0724) | Membrane protein | GAACATAGCTAGGTTGAGCCTCTCAACGC<br>AGGAAGACTCCGAAGCGCAAGAGGAGGAG<br>CTAAAGTGCACAGCCAACGGGATCTCAAG<br>CACCGCGTATAGTGCGGAAAACCAGAGGT<br>TTGCCTACTACCCAGCAATAGCCCTTCCA<br>TTTAGGGCAACTAGGGCCTCCGCCACGCA<br>GTGCGGGCCCTTCTACTGTCTAGACGTCG<br>ACCCTAACAAGGCCTTCGGCCAAGGGATC<br>CTAGTGAGAATAAACTACGAGGGGAGGAA<br>CATTTACTTCTACATAGGCTGGGTACCAG<br>CAGTTAAGGG |
| 25 | 580496 | 580795 | + | intragenic | SACI_RS03450 (Saci_0724) | Membrane protein | AACAAGGCCTTCGGCCAAGGGATCCTAGT<br>GAGAATAAACTACGAGGGGAGGAACATTT<br>ACTTCTACATAGGCTGGGTACCAGCAGTT<br>AAGGGATCGACTATCTTGGGAGTAGTCCA<br>GGGAGGTAAGTCCTTCAAGTCCTTATCCT<br>TGGCAAAGTACTTAGAGGAGTTCGGCATG<br>GATCTGAACACCGTGGGGGAAAGCTCGG<br>AGTAATCCCTGTTTACCCGCTGTCTAAGG<br>ACGTGGATGTGTGGTACAACCCGCCAATT<br>TACCAAGGCAACACAACTATCGCCTCAAT<br>AATACCTTGG |

| | | | | | | | |
|---|---|---|---|---|---|---|---|
| 26 | 1007000 | 1007299 | - | intragenic | SACI_RS05660 (Saci_1187) (*iunH*) | Nucleoside hydrolase | GCAATATCAACTATGGCGTCTAATGCCTT<br>TTTGTTCTCAGGCTTGAGTTTTTGTGGCT<br>TTGCAATTTTATCCCCTATTCCTCCTTTA<br>CCATGAACCTCTTCTACACTCCTAAAGTC<br>CTTGACCAAAGGCTTATCACTGCCAGGAT<br>AGACCTTAACATTTTCCTCTCCTATATAC<br>TCTAGAGCCCATAACGCATTATTAACCTC<br>TTGTTGGTAAGAAATGTTCCCCTCAACAA<br>TTGTTACCCCTTCCACACTCACTTTGTGT<br>CTCAAAAGCATGAACAAACTCATTATATC<br>GTCCTCTGCG |
| 27 | 1130248 | 1130547 | + | intragenic/ intergenic | SACI_RS06315 (Saci_1322) (*alba*) | Alba Nucleoid-associated protein (NAP) | TAGGTGGTTTAAGTGAGTTCAGGCGCAAC<br>CCCAAGCAATGTAGTATTGGTAGGTAAAA<br>AACCTGTAATGAACTATGTATTAGCAGCC<br>CTAACTCTACTAAATCAGGGTGTAAGTGA<br>GATAACAATAAAAGCTAGAGGTAGAGCAA<br>TAAGCAAAGCAGTTGACACTGTAGAAATA<br>GTGAGGAACAGATTCTTACCAGACAAAAT<br>AGAGGTAAAAGAGATTAGAATAGGAAGCC<br>AAGTTGTAACCAGCCAGGACGGAAGGCAG<br>TCTAGAGTTTCCACAATAGAAATAGGAAT<br>AAGAAAAAAG |
| 28 | 1309139 | 1309438 | + | intragenic/ intergenic | SACI_RS07310 (Saci_1534) | Thermostable acid protease | TATAACGCCTCTCCTGGAGAGCAGGTAAC<br>AATTAACGGCCCGATATACGAGAAACTCG<br>TAGTCCAGCTCAACTACACGTTCCTTGAG<br>GAGTTGGTTATACCCATTGTAACGGCTGT<br>CGTGATTGTTGTAGCGATAACGAGGAGGA<br>GATAGACAGTGAGTTAATGAATAGAAGTT<br>AGTACGCTAAACCCTTTTTCCTCGTTCTA<br>GAGTCTGCATAATACGAGAATACAAAGGC<br>AATAAACTGGAGATTTCAGAGGTGCCCTT<br>GGTCTCAAACGTTTATAACTTCTTGTTTA<br>ACCCATCACT |

| | | | | | | | |
|---|---|---|---|---|---|---|---|
| 29 | 1429094 | 1429393 | + | intergenic/ intragenic | SACI_RS07975 (Saci_1671) | Conserved hypothetical membrane protein | GTAAATTTTTAAATCATTTGGACTAAATA<br>TAATATAGAGAGAGTGAAATGACCAAGGT<br>TGAGGTGATTAAAATTGAGATCAGTGAGC<br>CATGCGGTCGTAAGCTGGACTCTCTGGAG<br>TCTAGCGCTATATCTCACACTTCCCTATC<br>TTTCTTCTGCCATCGAACACTCAATTAAT<br>AATCCACTTTTTGTGGGTCCCTTTTACAT<br>AATCTCCTCAGGTATTGGTAGTCTAAGTT<br>TATTAGCTCTTGAACGTCACGTGAAACTG<br>CTATCAACTATAGGGTTAGCACTTTCAGG<br>CATAGGACTG |
| 30 | 1486672 | 1486971 | - | intragenic/ intergenic | SACI_RS08250 (Saci_1727) | DUF1028 domain-containing protein | ACCTAATTGTCTTTTTTCTTTAAGCTGGT<br>CTGCAGAAGTTAAACTATTCACCGTACTC<br>CTTGCGTCATAACCCTTCTCTAGGAGTTC<br>TAATCCTTTTTTACCATACTCTAGATTAG<br>CTAGGGCTTGAGTTGCTATTGCCCCTACC<br>TCTGGTTTCAACCAAGGGACAAAAGCACC<br>CACTGCTAAGAACTTACTGGCAACACCTA<br>CTCCCCATGCTTCCTCGTTTGGGTCGTAA<br>ATTACTATTGAGTATGTCATATGTGTAAT<br>ATATACTACTGAAGGTATTTTATAATTAT<br>ATAATTCTCT |
| 31 | 1531730 | 1532029 | - | intragenic | SACI_RS08465 (Saci_1770) | Phenylacetic acid catabolic protein | TCATCAGAATTACGTTGTAGCTGTCAATT<br>ACGAAAACAAACGCCACGGCGTCCTCCCA<br>TGTAAATAAGGGCAAATTGAAGGGCTCTA<br>ACCTATGTAATCCAAGCCTCAGTTCTTGT<br>AGCTCTTTAACCTTATCCTTTTCACCAAA<br>GTCCTCGATCATACGAGACATCTGCCATG<br>CGTGATTCAGCTCATCTGCGACAAACCTA<br>GCAGTGAATAGTCTGGAGTCCACAGTTGG<br>GGAGTTAACTAGCCATGGAGATGTTTGCT<br>CCACAATAGCTAGTTTAGAATCGGCGATC<br>ACAAAAATTA |

| | | | | | | | |
|---|---|---|---|---|---|---|---|
| 32 | 2039264 | 2039563 | - | intragenic | SACI_RS10640 (Saci_2201) (*sir*) | Nitrite/sulfite reductase | CCCAAGTGTAAATCCCTTATTAGATTCAC<br>CTTAACTACTGGACCTAATTCCACACCAC<br>TATATTCCTTTATTTTCTCTCTCATCCAG<br>ATAATCCCAAATTTATGAACAACGTATTT<br>CAACCTAGCCCAATGCCTGTTCTTCCTAT<br>CCCCCCAATCTTGTTGTATCCTGACTATA<br>GAGTCTAAAACTTTCAATAGGTCGTCCTC<br>ACTTACTGTACCCAAAGGAAGTGCTAAAG<br>CAGAAAATGTCGGATAACCATTATTCTCG<br>CCCATTCCTCCACCAACATACAACTGATA<br>TTTCTCTATC |

**Table S2. SegB binding sites identified by ChIP-seq on *S. acidocaldarius* chromosome.** ChIP-seq enrichment peak sequences and relative genomic coordinates. Sites determined within the peak sequence through MEME are underlined and highlighted in yellow and those identified by DNase I footprint are shown in blue.
